# The NLRP3/eIF2 axis drives cell cycle progression in acute myeloid leukemia

**DOI:** 10.1101/2021.06.25.449862

**Authors:** Michela Luciano, Constantin Blöchl, Julia Vetter, Laura Urwanisch, Theresa Neuper, Dominik P. Elmer, Renate Bauer, Hieu-Hoa Dang, Helen Strandt, Daniel Neureiter, Peter Krenn, Suzana Tesanovic, Sebastian Rieser, Olivia Bergsleitner, Lukas Zell, Stephanie Binder, Susanne Schaller, Dirk Strunk, Lisa Pleyer, Richard Greil, Stephan Winkler, Tanja N. Hartmann, Christian G. Huber, Fritz Aberger, Jutta Horejs-Hoeck

**Author notes:** **Corresponding author:** Jutta Horejs-Hoeck, Department of Biosciences, Paris-Lodron University Salzburg Hellbrunner Strasse 34, 5020 Salzburg, Austria. **Key points** - NLRP3, ASC and IL1β are overexpressed in AML patients, with high expression of NLRP3 correlating with poor survival of AML patients. - Inhibition of NLRP3 reduces AML cell proliferation and cellular engraftment.

## Abstract

Aberrant activation of the NLR family pyrin domain containing 3 (NLRP3) inflammasome mediates numerous inflammatory diseases. Oncogenes can activate the NLRP3 inflammasome and thereby promote myeloproliferative neoplasia, suggesting a crucial role of NLRP3 in the malignant transformation of hematopoietic cells. Here, we show that bone marrow-derived mononuclear cells of AML patients display enhanced expression of NLRP3, IL-1β and IL-18 and that high-level expression of NLRP3 is linked to poor survival of AML patients. Pharmacological and genetic inhibition of NLRP3 inflammasome activation attenuated cell proliferation of MOLM-13 AML cells *in vitro*. *In vivo*, genetic inhibition of NLRP3 in MOLM-13 AML cells resulted in reduced engraftment potential in xenografts, along with reduced splenomegaly and organ infiltration. Differential proteomic analysis revealed the eIF2 pathway as potential target of NLRP3 in AML, with a significant increase of eIF2α phosphorylation upon NLRP3 inhibition. NLRP3 inhibition also caused a strong decrease in cyclin – dependent kinases CDK4 and CDK6, accompanied by an upregulation of the CDK inhibitor p21 (CDKN1A) and a marked arrest of cell cycle progression in the G0/G1 phase, consistent with the role of eIF2α phosphorylation as negative cell cycle regulator.

Taken together, we show that inhibition of the NLRP3 inflammasome reduces AML cell proliferation by promoting eIF2α phosphorylation, which in turn enhances the expression of cell cycle arrest genes such as p21. Thus, the study uncovers the NLRP3/eIF2 axis as new driver of AML proliferation and proposes a novel therapeutic treatment of AML by targeted inhibition of NLRP3 activation.

## Introduction

Chronic, non-resolving inflammation contributes critically to cancer development.^1^ In myeloid cells, inflammation is often triggered by inflammasomes, which are cytosolic multi-protein complexes that mediate the host defense against microbial infection and cellular stress. The NLRP3 inflammasome, one of the best-studied members of the inflammasome family,^2,3^ is composed of NLRP3 (NLR family pyrin domain containing 3), the adaptor protein ASC (Apoptosis-associated Speck-like protein containing CARD) and procaspase-1.^4^ Upon inflammasome formation, procaspase-1 undergoes proteolytic cleavage and the resulting active caspase-1 converts the cytokine precursors pro-IL-1β and pro-IL-18 into mature IL-1β and IL-18 which induce inflammation and pyroptotic cell death.^5^ NLRP3 also seems to play a role in the development and expansion of hematopoietic stem progenitor cells (HSPCs) as well as in hematologic pathologies.^6^ For example, in myelodysplastic syndrome (MDS), aberrant activation of the NLPR3 inflammasome leads to the release of IL-1β and IL-18, resulting in pyroptotic cell death. Accordingly, knockdown or pharmacological inhibition of NLRP3 was shown to restore effective hematopoiesis in bone marrow (BM)-derived mononuclear cells isolated from MDS patients.^7^ The potential role of the NLRP3 inflammasome as a driver of myeloid malignancies was further supported by the finding that oncogenic *Kras* mutation is functionally linked to activation of NLRP3, implicating activation of the RAC1/ROS/NLRP3/IL-1β axis as a crucial event in myeloproliferative disorders.^8^ Although IL-1β is a known growth factor for acute myeloid leukemia (AML) cells *in vitro*,^9^ and several hematopoietic cancers including acute and chronic leukemias show constitutive production of high IL-1β,^10^ the role of IL-1β in human AML remains controversial and data on the role of inflammasomes as promoters of innate inflammation in leukemias are scarce. Here, we investigated the role of NLRP3 in human AML and show that inhibition of NLRP3 results in phosphorylation of the translation initiation factor eIF2α, downregulation of the cell cycle proteins CDK4 and CDK6, and reduced proliferation of AML cells.

## Methods

### Primary human AML samples, cell lines and culture conditions

All studies involving human cells were conducted in accordance with the guidelines of the World Medical Association’s Declaration of Helsinki. Following written informed consent, BM aspirate samples were collected from patients with newly diagnosed AML (Ethics committee Salzburg approval: 415-E/2009/2-2016). BM from healthy donors was obtained from Caltag Medsystems, UK (www.caltagmedsystems.co.uk). Human BM mononuclear cells, CD34^+^ umbilical cord and CD34^+^ hematopoietic stem/progenitor cell (Ethics Committee Salzburg approval 415-E/1776/4-2014) were isolated using density gradient centrifugation (Lymphoprep; Stemcell Technologies). Primary human cells, human AML cell lines HL-60, MOLM-13, MV4-11 and the human cervix carcinoma cell line (HeLa) were cultured in RPMI 1640 medium or Minimum Essential Medium Eagle (MEM) (HeLa) supplemented with 10% heat-inactivated FBS, 1% penicillin and 1% streptomycin and non-essential amino acids (HeLa). For NLRP3 inhibition, the small molecule CP-456773 sodium salt ≥98% (HPLC) (Sigma) was used.

### Generation of NLRP3-deficient MOLM-13 cells (MOLM-13 ΔNLRP3)

To generate control or NLRP3 knockout (ΔNLRP3) MOLM-13 cells, lentiviral particles (0.5 MOI) produced in HEK293FT cells transiently transfected with two packaging plasmids (psPAX2 and pMD2-G) and the lentiCRISPR v2 plasmid (Addgene #52961)^11^ containing a non-targeting (GGCATCTTAACTAATCGTCT) or NLRP3-targeting (AAAAGAGATGAGCCGAAGTG; CRISPick design tool^12,13^) sgRNA were used to transduce MOLM-13 cells via spin-infection.^14^ Successfully transduced cells were selected for puromycin resistance. NLRP3 protein knockout validation was performed by Western blot analysis.

### Immunohistochemical staining and detection of cytokines

Immunohistochemistry (IHC) for NLRP3/NALP3 (Adipogen Life Sciences, AG-20-b-0014-C100, Liestal, Swiss) was performed on prepared cell blocks of the human cancer cell lines MOLM-13 and MOLM-13 ΔNLRP3 as well as on routinely archived formalin-fixed paraffin-embedded (FFPE) BM trephine samples of normal controls (n=10) and AML cases (n=13, primary diagnosis n=11) (AML-M5 NOS according to WHO classification). Sections were stained with primary NLRP3/NALP3 antibody using a Benchmark Ultra platform (Ventana, Tucson, USA) and an ultraView Universal DAB Detection Kit (Ventana) after application of the amplification kit (Ventana). The results were scored by assessing the extensity (% positive cells) and intensity of IHC staining (0-3) on three different representative microscope fields and expressed semi-quantitatively using the quickscore method by multiplication of the extensity and intensity (yielding values between 0-300) for each field.^15^ Analysis of cytokine secretion in primary human AML samples was performed using the Cytokine/Chemokine/Growth Factor 45-Plex Human ProcartaPlex™ system (ThermoFisher).

### Cell proliferation and cell cycle analyses

For proliferation analyses, 1×10^7^ cells were stained with the proliferation dye (eBioscience™ Cell Proliferation Dye eFluor™ 450, Invitrogen™) diluted 1:5000 in PBS and cultured for 72 h. After harvesting, the cells were counted using a Neubauer chamber and proliferation was assayed by flow cytometry on a BD FACS Canto II (BD Biosciences). Cell viability was analyzed using the fixable viability dye eFluor™ 780 (eBioscience™). Cell cycle analysis was performed using the FxCycle™ PI/RNase Staining Solution according to the manufacturer’s instructions (Invitrogen, Molecular Probes by Life Technologies, Catalog number: F10797).

### qRT-PCR

Total RNA of cultured cells was extracted with Trizol (Tri Reagent^®^, Sigma) according to the manufacturer’s instructions. Complementary DNA (cDNA) was generated using RevertAid H Minus M-MulV reverse transcriptase (Thermo Fisher Scientific). Expression levels were determined by quantitative real-time PCR on a Rotorgene 3000 (Qiagen Instruments, Hombrechtikon, Switzerland) using Luna^®^ Universal Probe qPCR Master Mix (New England BioLabs^®^ Inc). The large ribosomal protein P0 (RPLP0) was used as a reference gene. Relative mRNA expression (x) was calculated x = 2^−ΔCt^, where Δct represents the difference between the threshold cycle (ct) of the NLRP3 gene and the reference gene. NLRP3 primer pair (Sigma): forward 5’ -TCAGCACTAATCAGAATCTCACGCACCTTT -3’ and reverse 5’ -CCAGGTCATTGTTGCCCAGGCTC -3’; RPLP0 primer pair: forward 5’ – GGCACCATTGAAATCCTGAGTGATGTG -3’ and reverse 5’ -TTGCGGACACCCTCCAGGAAG -3’.

### Western blot

Pellets of cultured cells were harvested and lysed in 80 μL of 2x Laemmli sample buffer (BIO-RAD) with 5% β-mercaptoethanol (Sigma-Aldrich). Samples were separated on 4% to 12% gradient gels (NuPAGE, Life Technologies) and subsequently blotted onto a nitrocellulose membrane (BIO-RAD) using a semi-dry transfer system. The following antibodies were used according to the manufacturers’ instructions: NLRP3 (D4D8T) (Cell Signaling), ASC (O93E9) (BioLegend), eIF2α (D7D3) XP^®^ (Cell Signaling), Phospho eiF2α (Ser51) (D9G8) XP^®^ (Cell Signaling), CDK4 (D9G3E) (Cell Signaling), CDK6 (DCS83) (Cell Signaling), p21 (12D1) (Cell Signaling), β-actin (Cell Signaling) and HRP-linked anti-rabbit secondary antibody (Cell Signaling). Western blots were quantified by measuring protein band intensity using ImageJ (NIH) software.

### Animal studies

12- to 16-week-old male NOD-scid IL2Rgnull-3/GM/SF (NSG-S) mice obtained from the Jackson Laboratories were kept under specific pathogen free conditions, in a 12-hour light-dark cycle, with a standard chow diet and water ad libitum at the animal facility of the University of Salzburg. All mouse experiments were approved by the Federal Ministry of Education, Science and Research (BMBWF), Austria (permission number: BMWFW-66.012/0032-WF/V/3b/2017) and complied with EU guidelines (2010/63/EU) and Austrian law (TVG 2012).

For the MOLM-13 tumor engraftment studies, NSG-S mice were intravenously injected with 0.5 × 10^6^ MOLM-13 control cells or ΔNLRP3 cells and constantly monitored and scored for leukemia-associated changes in physical appearance, breathing rate and behavior. The experiment was repeated twice. All mice of the same experiment were euthanized when predefined termination criteria were met, which was at day 14 after injection in the first, and on day 19 in the second experiment. Peripheral blood, spleen and bone marrow were isolated and evaluated for the percentage of human CD45^+^ cells by flow cytometry (CD45-PerCP-Cy5.5, Biolegend Cat: 304028; BD FACS Canto II). Populations were defined and analyzed using FlowJo software (Flowjo v10.7.1, BD Biosciences).

### Proteomics

Cellular proteomes of CP-456773-treated MOLM-13 cells (0, 75 and 125 μg/mL) were prepared utilizing S-Trap columns (Protifi, USA) with small amendments to the manufacturer’s instructions, as detailed in the supplemental Methods. Peptides were labeled by TMT-10plex™ (Thermo Fisher, USA) followed by concatenated off-line high-pH reversed phase fractionation of multiplexed samples.^16,17^ Fractions were separated on an Thermo Scientific™ UltiMate™ 3000 RSLCnano system (Thermo Fisher Scientific, Germering, Germany) equipped with a 2000 mm μPAC™ column (PharmaFluidics, Belgium) coupled to a benchtop quadrupole-Orbitrap instrument (Thermo Scientific™ Q Exactive™ Plus). Instrument parameters are supplied in the supplemental Methods. Acquired data were evaluated using MaxQuant (v1.6.3.4) to provide Uniprot entries and further processed in Perseus (v1.6.6.1).^18–20^ Significantly regulated proteins were investigated in Ingenuity Pathway Analysis (v47547484, Qiagen, USA)

### Data availability

The mass spectrometry proteomics data have been deposited in the ProteomeXchange Consortium via the PRIDE partner repository with the dataset identifier PXD026293.^21^

### Database analysis

Public genome datasets GSE13159 and GSE12417 from NCBI’s Gene Expression Omnibus (NCBI-GEO) were used. Based on the GSE13159 dataset, IL-1β, IL-18 and NLRP3 expression was evaluated in AML patients and in healthy individuals. The dataset used is part of the MILE (Microarray Innovations in Leukemia) study research program. The study includes whole-genome analysis data from 542 AML patients, 76 CML patients, and 74 healthy donors for a total sample size of 2096 from 11 participating centers on three continents.^22,23^ The GSE12417 dataset was used to analyze NLRP3 expression among AML French-American-British (FAB) subclasses.^24^ The GSE12417 dataset includes gene expression profiling data of 163 patients treated in the German AMLCG 1999 trial to predict the overall survival (OS) in cytogenetically normal AML (CN-AML).^25^ Datasets from the GEO database were imported using GEOparse and the analysis was performed using Python.

### Statistical analysis

Statistical analyses were performed with GraphPad Prism 8 software (Graphpad Software, San Diego, CA, USA). Comparisons in multiple groups were analyzed by one-way ANOVA including a post-hoc test. For comparison between two groups, paired Student’s t-test was used. p values <0.05 were considered to be significant.

## Results

### Increased expression of the NLRP3 inflammasome components NLRP3, ASC, IL-1β and IL-18 in bone marrow samples of AML patients

To investigate whether components of the NLRP3 inflammasome are aberrantly expressed in BM of AML patients, we analyzed the public genomic dataset GSE13159 from NCBI’s Gene Expression Omnibus (NCBI-GEO),^26^ comprising whole-genome data from 542 AML patients and 74 healthy controls.^22,23^ Figure 1A shows that NLRP3 and the downstream targets ASC, IL-1β and IL-18 were significantly upregulated in AML patients compared to healthy controls, whereas genes encoding other members of the NLRP protein family involved in inflammasome formation were not affected (supplemental Figure 1A). Enhanced NLRP3, IL-1β and IL-18 levels were confirmed at the protein level by immunohistochemistry and by multiplex analyses applied to bone marrow mononuclear cells (BM-MNCs) derived from our own cohort of AML patients (Figure 1B-D). Evaluation of the outcome of AML patients with high (n=40) or low (n= 39) NLRP3 expression data from the GEO:GSE12417 (CN-AML) series assessed by Log rank analysis of survival curves revealed a significantly prolonged median OS among all patients with low NLRP3 expression compared to those with high NLRP3 expression (Figure 1E). This indicates that NLRP3 may indeed play a crucial role in the pathogenicity of AML. As mice carrying an inducible Kras^G12D^ mutation show enhanced NLRP3 expression,^8^ we determined associations between NLRP3 components and oncogenic kinases in BM of AML patients by calculating correlations using the GSE12417 dataset. We found correlations between NLRP3 and each of NF-κB1 (R=0.403), NRAS (R=0.449), FLT3 (R=0.299) and KRAS (R=0.220) (supplemental Figure 1B). Of note, NLRP3 and ASC were highly expressed in patients with acute monoblastic/monocytic leukemia (FAB M5) (Figure 1F). Accordingly, MOLM-13 and MV4-11, two FLT3-ITD mutated cell lines representing this FAB subtype, show high NLRP3 expression at the protein level, compared to the FAB M2 cell line HL-60, even though the latter cell line carries an NRAS mutation. In addition, we identified the cervical cancer cell line HeLa as NLRP3 negative and thus used this cell line as a negative control for subsequent functional experiments targeting NLRP3 activity (Figure 1G).

**Figure 1.**
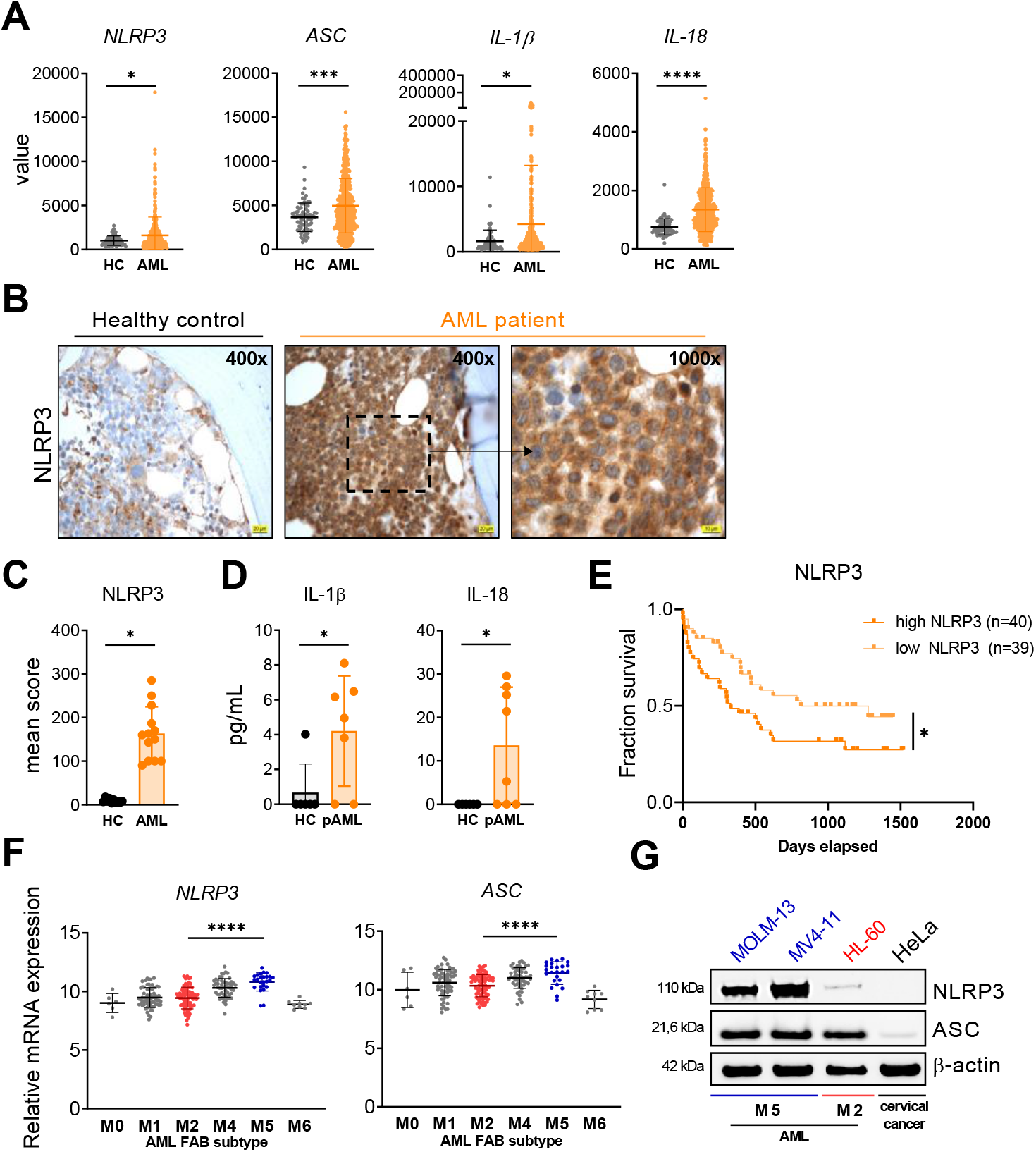
Increased NLRP3 and ASC expression in AML patients. (A) NLRP3, ASC, IL-18 and IL-1β expression in AML patients (AML, n=542) and healthy individuals (HC, n=74) was determined using a publicly available dataset GSE13159, which was analyzed using Python. (B-C) Representative immunohistochemistry NLRP3 staining (B) and mean score of FFPE BM trephine samples (C) of normal controls (HC, n=10) and AML cases (AML, n=13) are shown. (D) Detection of IL-1β and IL-18 secretion using Multiplex Assay in BM-MNCs isolated from AML patient samples (AML, n=8) healthy controls (HC, CD34^+^ hematopoietic stem/progenitor cells G-CSF immuno-mobilized, and umbilical cord blood derived CD34^+^ stem/progenitor cells, n=6) cultured for 24 hours. (E) Kaplan-Meier curve analysis of the dataset GSE12417 (platform GPL570) comparing the survival of AML patients with high NLRP3 expression (n=39) vs low NLRP3 expression (n=40) was performed using Python. A two-tailed, unpaired t test was used for the statistical analysis between two groups. Dots represent individual donors, mean ± SD are shown. *p ≤ 0.05, ***p ≤ 0.001, ****p ≤ 0.0001. (F) NLRP3 and ASC expression analysis within the AML subgroups was performed using the public genome dataset GSE12417. Datasets from the GEO database were imported using GEOparse and the analysis was performed using Python (AML FAB 0, n =6; AML FAB 1, n=68; AML FAB 2, n=79; AML FAB 4, n=53; AML FAB 5, n=25; AML FAB 6, n=9). Single dots indicate individual donors, the horizontal lines in each column define mean ± SD. One-way ANOVA with Tukey’s post-hoc test was performed. **** p<0.0001 (G) NLRP3 and ASC expression was detected in the human AML cell lines MOLM-13 and MV4-11 (both AML FAB subtype 5), HL-60 (AML FAB subtype 2) and cervical cancer cells (HeLa) by Western Blot. One representative immunoblot out of 3 is shown.

### NLRP3 inhibition results in reduced cell proliferation

To analyze whether inhibition of NLRP3 inflammasome activation has any effect on AML cell proliferation, we applied the NLRP3-specific inflammasome inhibitor CP-456773 (also referred to as CRID3 sodium salt or MCC950) to MOLM-13 cells. CP-456773 has previously been shown to specifically block NLRP3-induced ASC oligomerization while not affecting other inflammasomes.^27^ Accordingly, IL-1β secretion, induced upon activation of the NLRP3 inflammasome by LPS and ATP ^27,28^ was significantly blocked by CP-456773 treatment (supplemental Figure 2A). To test the effect of CP-456773 on AML cell proliferation, we cultured NLRP3-positive MOLM-13 and for comparison NLRP3-negative HeLa cells (Figure 2A) for 72 hours in the presence of increasing CP-456773 concentrations. As shown in Figure 2B, CP-456773 treatment of MOLM-13 resulted in a clear and concentration-dependent reduction in cell numbers, which was replicated in MV4-11 and HL-60 AML cells (supplemental Figure 2B). In contrast, the inhibitor had no effect on the proliferation of NLRP3-negative HeLa cells (Figure 2B/C), clearly showing that CP-456773 selectively acts on cells expressing NLRP3. To further support the crucial role of NLRP3 in AML, we generated NLRP3-deficient MOLM-13 cells (ΔNLRP3) by means of CRISPR/Cas9 technology (Figure 2D/E). In line with the pharmacological inhibition of NLRP3, genetic inactivation of NLRP3 in MOLM-13 cells (ΔNLRP3) resulted in decreased proliferation compared to MOLM-13 proficient cells (Figure 2F).

**Figure 2.**
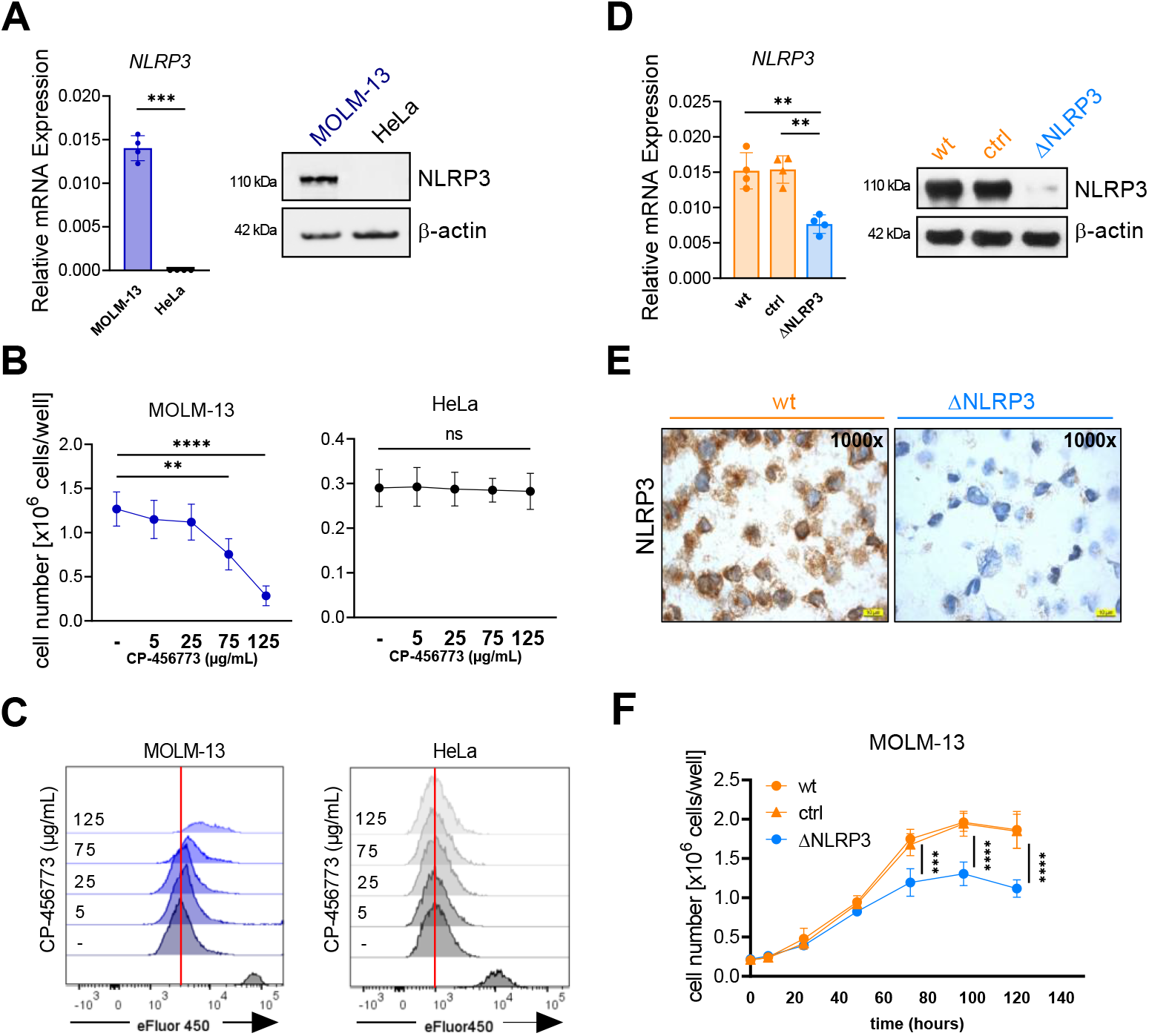
Decreased cell proliferation upon NLRP3 inhibition with CP-456773 and NLRP3 deletion in MOLM-13 cells. (A) NLRP3 mRNA and protein expression was determined in MOLM-13 and HeLa cells by qRT-PCR and Western Blot. One representative immunoblot out of 3 is shown. (B) MOLM-13 and HeLa cell numbers upon treatment with the indicated concentrations of CP-456773 for 72 hours were evaluated using a Neubauer chamber (MOLM-13, n=5; HeLa, n=4). (C) Proliferation of MOLM-13 and HeLa cells in the presence of the indicated concentrations of CP-456673 was monitored after 72 hours by flow cytometry using the Cell Proliferation Dye eFluor 450. Histograms of one representative out of 5 for MOLM-13 and 4 for HeLa experiments are shown. The vertical red lines indicate the fluorescence peak of proliferating cells in the absence of CP-456773. Freshly stained MOLM-13 served as negative control (bottom peak). (D) Analysis of NLRP3 mRNA and protein expression in MOLM-13 wild-type (wt), MOLM-13 CRISPR/Cas9 (ctrl) and MOLM-13 CRISPR/Cas9 NLRP3 knock-out (ΔNLRP3) cells by qRT-PCR and Western Blot. One representative immunoblot out of 3 is shown. (E) Immunohistochemical staining of NLRP3 performed in wt and ΔNLRP3 MOLM-13 cells. (F) MOLM-13 wt, ctrl and ΔNLRP3 cells were harvested at the indicated timepoints and counted with a Neubauer chamber (n=6). A two-tailed, unpaired t test was performed for the statistical analysis between two groups, while one-way ANOVA with Tukey’s post-hoc test was performed for multiple comparisons. Data represent mean ± SD. ***p ≤ 0.001, ****p ≤ 0.0001.

### Decreased engraftment of MOLM13 ΔNLRP3 cells

To evaluate the consequences of NLRP3 deletion *in vivo*, immunodeficient NSGS mice (see materials and methods) were engrafted with MOLM-13 control (ctrl) or MOLM-13 ΔNLRP3 cells lacking NLRP3 and sacrificed when the degree of distress was evaluated as moderate according to our scoring protocol (Figure 3A). NSGS mice injected with PBS only, were used as additional control and healthy reference cohort. Compared to PBS-injected mice, mice engrafted with NLRP3-positive MOLM-13 ctrl cells showed a significant loss of body weight and developed severe splenomegaly, while MOLM-13 ΔNLRP3 did not suffer from body weight loss and showed significantly reduced splenomegaly (Figure 3B/C). In addition, recipient mice engrafted with MOLM-13 ΔNLRP3 cells showed reduced percentages of total human CD45^+^ cells in peripheral blood, spleen and bone marrow, compared to mice engrafted with MOLM-13 ctrl cells (Figure 3D). These data indicate that NLRP3 deficiency in MOLM-13 cells reduces engraftment and disease progression of AML *in vivo*.

**Figure 3.**
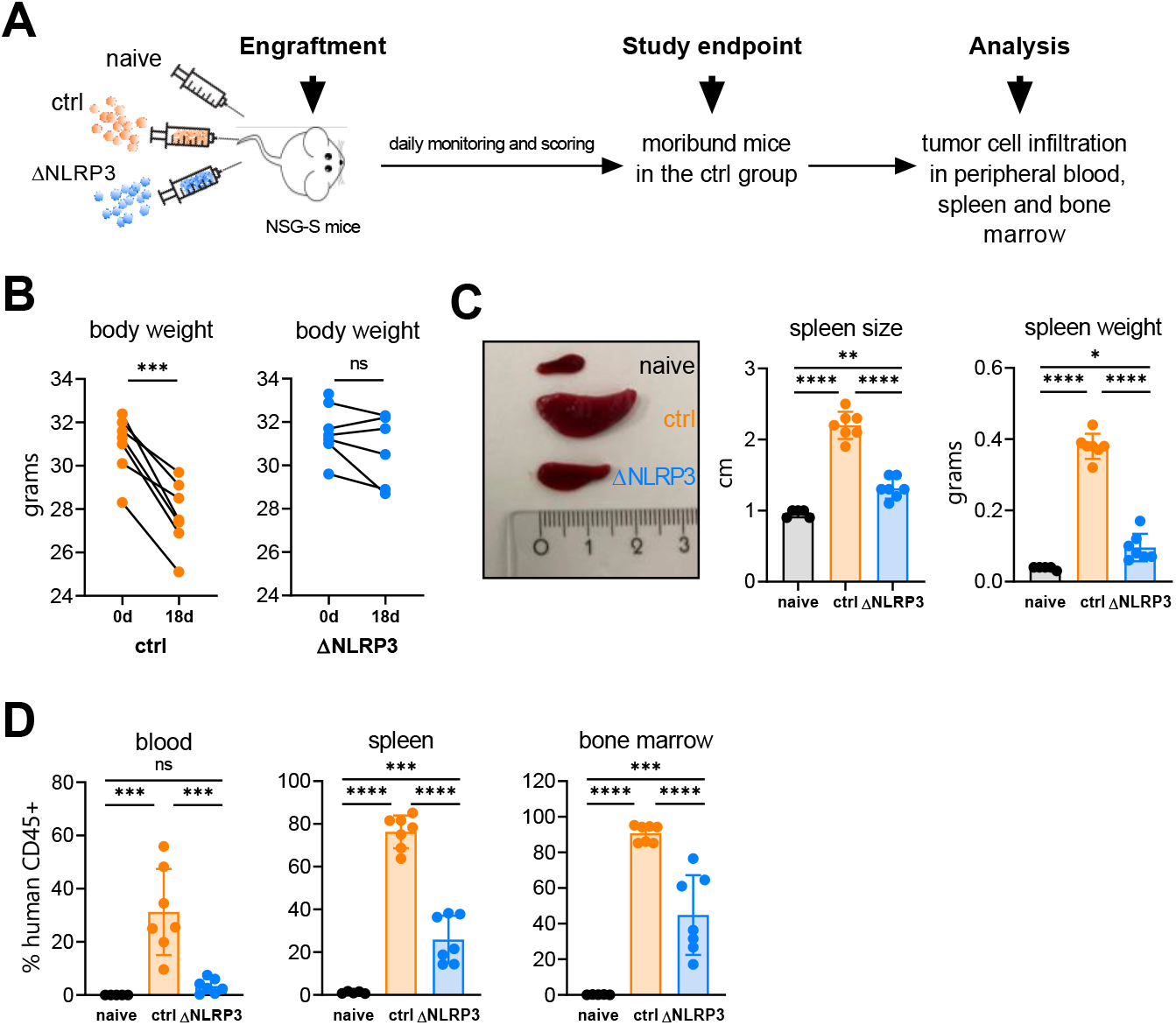
NLRP3 promotes AML development and progression *in vivo*. NSG-S mice receiving PBS (naïve), CRISPR/Cas9 control MOLM-13 cells (ctrl), or CRISPR/Cas9 NLRP3 knockout MOLM-13 cells (ΔNLRP3) were monitored for leukemia development and analyzed for tumor cell infiltration. (B) The study endpoint was defined as the time point when the first ctrl mice became sick and moribund, best reflected by their rapid weight loss. (C) Engraftment parameters spleen length and weight (C) and human CD45+ cells in peripheral blood, spleen and bone marrow measured by flow cytometry (D) at the study endpoint. Statistical analyses were performed by unpaired t-test, two-sided or one-way ANOVA, Tukey’s multiple comparison test (C,D). n = 5 for naïve samples, n = 7 for ctrl samples, and n = 7 for ΔNLRP3 samples. Data in panels C and D are shown as mean ± SD. *p ≤ 0.05, **p ≤ 0.01, ***p ≤ 0.001, ****p ≤ 0.0001.

### Impact of NLRP3 inhibition on eIF2 signaling

To evaluate the molecular mechanisms mediating the effects of NLRP3 as a driver of AML cell proliferation, MOLM-13 wild-type cells were treated with CP-456773 (75 and 125 mg/mL) and subjected to differential proteomic analyses using high-performance liquid chromatography (HPLC) employing micro pillar array columns (μPAC) in combination with Orbitrap mass spectrometry (MS) (Figure 4A). In total, 6950 proteins were identified, quantified and, according to global changes in protein expression patterns, assigned to the three different treatment groups based on multivariate statistics employing principal component analysis (PCA) (supplemental Figure 3A). In order to reveal significantly enriched proteins, we assessed adjusted p-values, correcting for multiple comparison by Benjamini-Hochberg with a false discovery rate of 10%. In addition, a Venn diagram was used to depict the overlap (orange) between significantly regulated proteins after treatment with 75 μg/mL CP-456773 (yellow) and 125 μg/mL CP-456773 (red) (supplemental Figure 3B). To obtain an overview of the regulated signaling pathways, significantly regulated proteins and their mean log2 fold changes of three replicates were used as inputs for pathway analysis. The regulated pathways were ranked according to their −log10 (p-value) by applying Fisher’s exact test, which revealed the eIF2 pathway as being the most affected by NLRP3 inhibition (Figure 4B). EIF2 is a key regulator of global protein synthesis in eukaryotic cells.^29^ It is well established that phosphorylation of eIF2α leads to blockade of global protein synthesis,^30^ but can also induce cell cycle arrest^31^ and apoptosis (Figure 4C).^32^ Thus, we investigated if NLRP3 deletion or inhibition of NLRP3 activation has an impact on eIF2α phosphorylation. Immuno-blotting shows enhanced phosphorylation of eIF2α both in the absence of NLRP3 expression (MOLM-13 ΔNLRP3) and upon treatment with the NLRP3 inhibitor CP-456773 (Figure 4D). The clear negative correlation between NLRP3 activation and eIF2α phosphorylation indicates that high NLRP3 activation, as observed in AML, might prevent eIF2α phosphorylation. In light of the regulatory role of eIF2, we therefore suggest that NLRP3 prevents eIF2-regulated cell cycle arrest and/or apoptosis and instead promotes the proliferation of leukemic cells.

**Figure 4.**
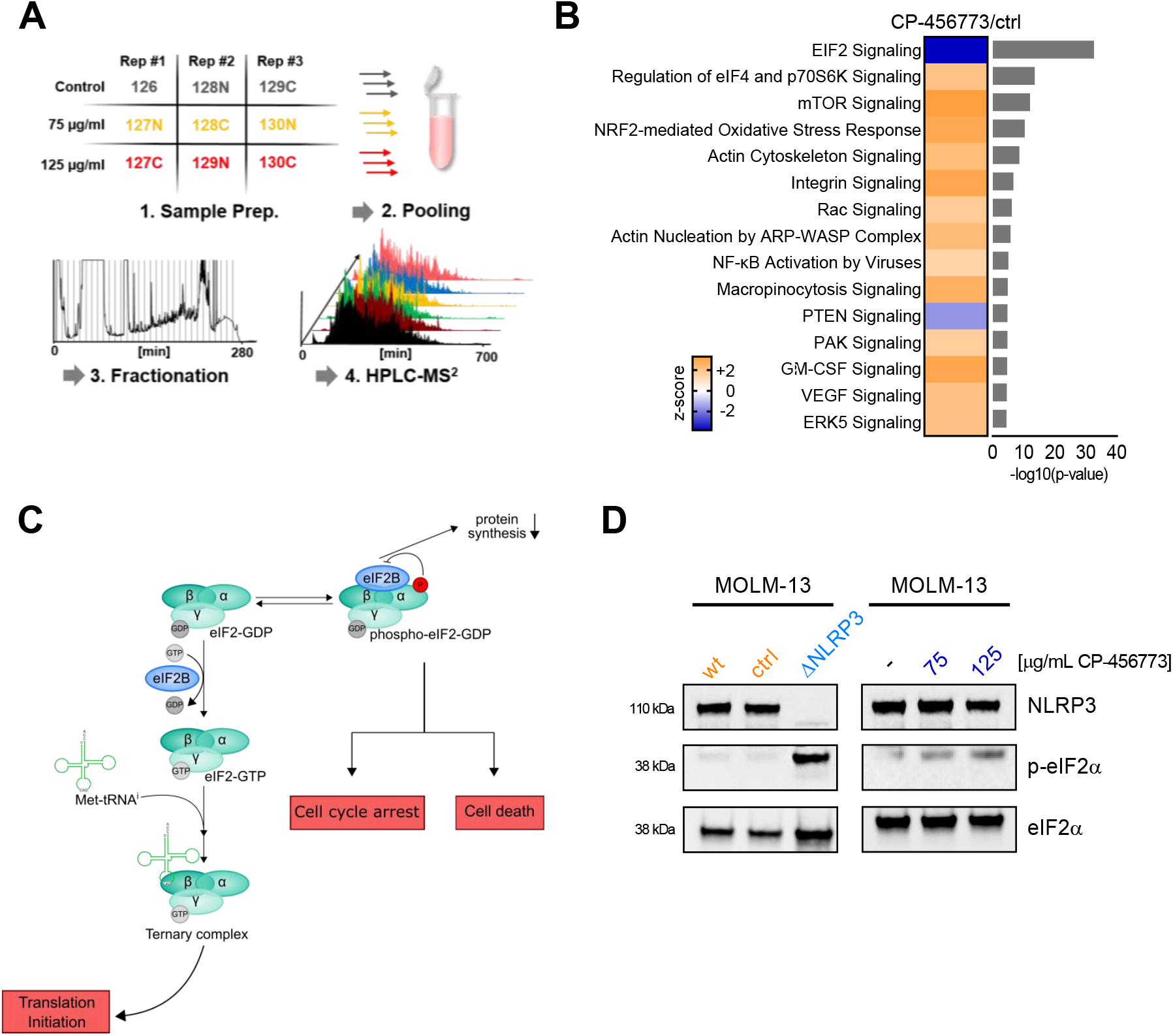
In-depth quantitative proteomics analysis of MOLM-13 cells treated with CP-456773 reveals eIF2α signaling pathway involvement. (A) Experimental set-up for the proteomics experiment. Cells were treated with either 0, 75 or 125 μg/mL of CP-456773 each in three biological replicates. After parallel proteomics sample preparation, peptides were labeled by tandem mass tags (TMT 126-130C) for relative quantitation of protein abundances and, eventually, were pooled. The combined sample was subjected to off-line high-pH reversed phase fractionation and subsequently analyzed by second dimension HPLC-MS^2^. (B) Overview of significantly altered canonical pathways. Pathways were filtered using a −log10(p-value) cut-off of 4.5 and a z-score threshold of ±1.5 and are ranked according to their −log10(p-value). The color code represents the z-score of a specific pathway whereas the corresponding bar graph on the right shows the respective log10(p-value). Depicted p-values of affected pathways were obtained from a Fisher’s exact test. (**C**) Graphic illustration of eIF2 signaling. Unphosphorylated eIF2α initiates protein translation in a GTP-dependent manner, whereas phosphorylated eIF2α favors cell cycle arrest and/or cell death. (D)Analyses of phosphorylated eIF2α in relation to NLRP3 expression or activation. MOLM-13 wt, ctrl and ΔNLRP3 cells (left panel) and MOLM-13 wt cells in the absence or presence of the indicated concentrations of CP-456773 (right panel) were cultured for 24 hours. Protein expression and phosphorylation were determined by Western Blot analysis. One representative immunoblot out of 3 is shown.

### NLRP3 inhibition with CP-456773 induces cell cycle arrest of MOLM-13 cells

As shown in the volcano plot (Figure 5A), important regulators of cell cycle progression were found to be differentially expressed between untreated cells and those treated with 125 μg/mL CP-456773. Cell cycle progression is a tightly regulated process governed by cyclin-dependent kinases (CDKs), which interact with cyclin-dependent kinase inhibitors (CDKIs) by forming stable complexes with CDKs before they are bound to cyclins (Figure 5B).^33^ Detailed protein expression analysis by quantitative proteomics and by immunoblotting showed that CDK6 and CDK4, two kinases promoting cell cycle progression, were significantly downregulated in presence of increasing CP-456773 concentrations, while the CDK inhibitor CDKN1A (also known as p21 or p21WAF1/Cip1) was significantly induced (Figure 5C-E). In line with these observations, cell cycle analysis by flow cytometry revealed that treatment with CP-456773 for 48 h increased the proportion of cells in G0/G1 phase at the expense of cells in S and G2/M phase, suggesting that pharmacologic inhibition of NLRP3 induces G0/G1 cell cycle arrest of AML cells (Figure 5F/G). Therefore, we propose that inhibition of NLRP3 promotes eIF2α phosphorylation, which in turn results in reduced AML cell proliferation and G1/G0 cell cycle arrest through the modulation of critical cell cycle regulators such as CDKN1A and CDK4/6.

**Figure 5.**
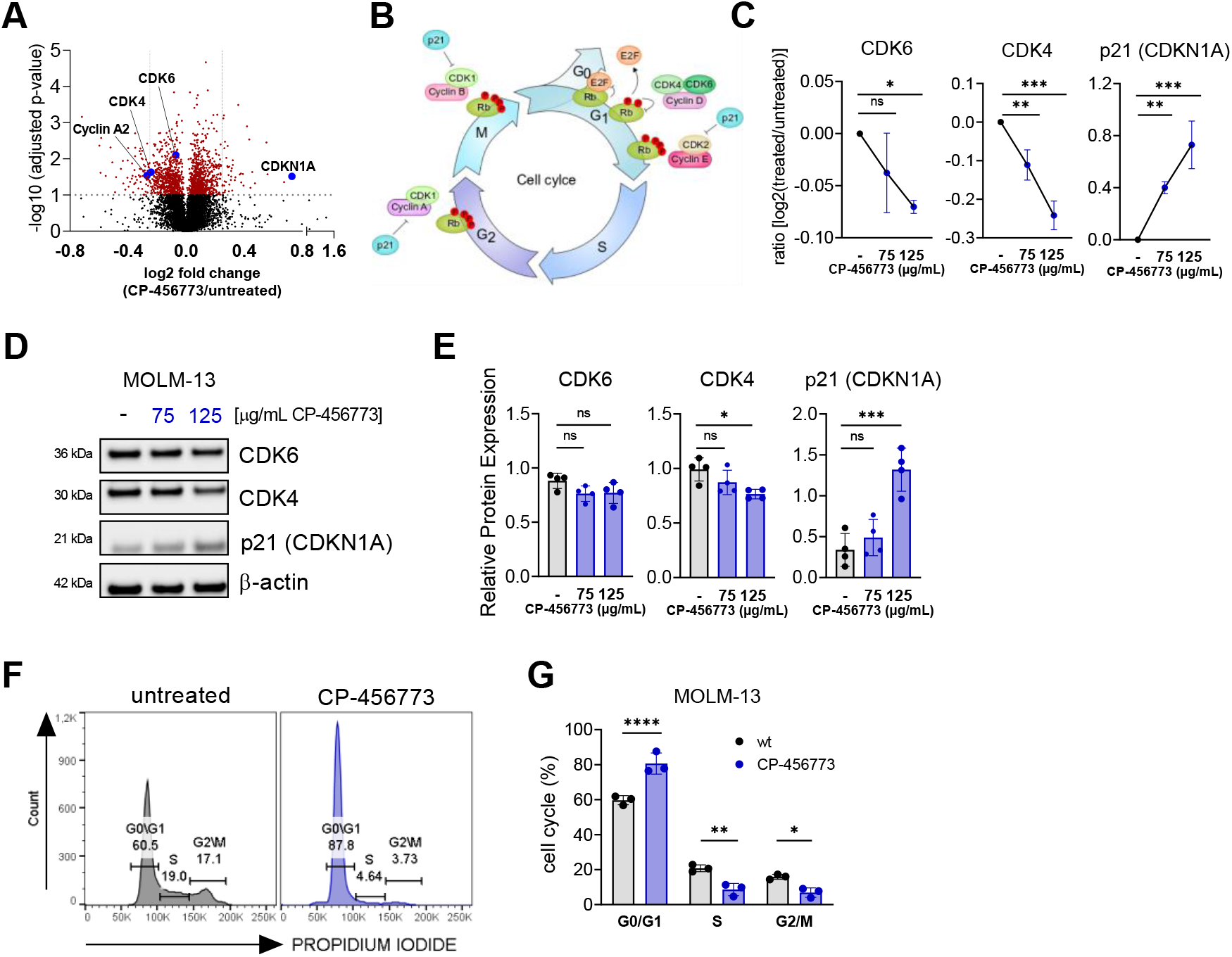
NLRP3 inhibition induces cell cycle arrest. (A) Volcano plot illustrating the differences (log2 FC) in protein expression between untreated MOLM-13 cells and MOLM-13 cells treated with CP-456773 (125μg/mL). Adjusted p-values were obtained from a paired t-test applying Benjamini-Hochberg correction. The horizontal gray dotted line indicates an adjusted p-value of 0.1 separating the significantly regulated proteins (red) from the rest (black) with an FDR of 10%. Regulated candidate proteins are showed as blue dots and labeled with their gene names. (B) Graphic illustration of cell cycle regulation. Cyclin-dependent kinase 4 (CDK4) and CDK6 are key drivers of the cell cycle involved in the progression from G0/G1 to S phase. The CDK inhibitor p21 (also known as p21(WAF1/Cip1 or CDKN1A) promotes cell cycle arrest at different stages of the cell cycle. (C) Protein expression values of CDK6, CDK4 and CDKN1A obtained by quantitative proteomics. The log2 fold changes in protein expression were plotted against increasing amounts of CP-456773. Statistical significance was tested by employing one-way ANOVA followed by Dunnett’s post-hoc test. *p ≤ 0.05, **p ≤ 0.01, ***p ≤ 0.001. Mean values (n=3) were used for plotting the data, and standard deviations are indicated. (D) Analysis of expression of the cell cycle regulators CDK6, CDK4 and p21 (CDKN1A) upon NLRP3 inhibition. MOLM-13 cells treated with CP-456773 (75 and 125 μg/mL) or left untreated were cultured for 24 hours and protein expression was determined by Western Blot. One representative immunoblot out of 5 is shown (E) Quantification of protein levels of CDK4, CDK6 and p21 (CDKN1A) from Western Blots (n=5) performed with ImageJ and normalized to β-actin. Dots indicate individual experiments; bars represent mean±SD. For statistical analysis a one-way ANOVA followed by Dunnett’s post-hoc test was used.*p ≤ 0.05, **p ≤ 0.01, ***p ≤ 0.001 (F) Cell cycle analysis of MOLM-13 wt and MOLM-13 wt treated with CP-456773 (125 μg/mL) cultured for 48 hours. Propidium iodide staining was used to assess the cell cycle state in each condition and analyzed by flow cytometry. Histograms of one representative experiment out of 3 are shown. (G) Bar diagram showing cell distribution in subG1, G0/G1, S and G2/M phases under the different conditions. Dots indicate individual experiments; bars represent mean±SD. For statistical analysis a two-tailed, unpaired t test was performed for the statistical analysis between two groups.*p ≤ 0.05, **p ≤ 0.01, ***p ≤ 0.001, ****p ≤ 0.0001.

## Discussion

While unbalanced release of pro-inflammatory cytokines such as IL-1, IL-6, TNF-α and GM-CSF has been frequently reported in AML,^34^ a leukemogenic role for the NLRP3 inflammasome — the central hub of innate immunity mediating the release of pro-inflammatory factors — has only recently been suggested.^8,35,36^

Important evidence for the role of NLRP3 in human cancers was provided by a study showing that 15 out of 24 different, mostly solid, cancers had significantly different expression profiles of NLRP3 and inflammasome-related genes compared to normal tissue samples.^37^ It has also emerged that NLRP3 can play dual roles in cancer, being either tumor suppressive, as observed in colitis-associated colorectal cancer (CAC), or tumor promoting, especially evident in cancers of the skin, breast and stomach.^38,39^ The role of NLRP3 in hematopoietic malignancies, however, is less clear. In the present study, we demonstrated that expression of NLRP3 and the NLRP3-inflammasome-related genes ASC, IL-1β and IL-18 is increased in bone marrow cells of AML patients compared to cells from healthy donors. Accordingly, Hamarsheh et al. reported that increased NLRP3 activity was associated with cytopenia, splenomegaly, and myeloproliferation, all features commonly observed in patients with chronic myelomonocytic leukemia (CMML), juvenile myelomonocytic leukemia (JMML), and AML.^8^ The latter study further describes oncogenic KRAS signaling as a driver of NLRP3 activation, and reports particularly high levels of cleaved caspase-1 and increased IL-1β production in peripheral blood mononuclear cells from KRAS^mut^ AML patients compared to AML patients without KRAS mutations.^8^ Our study extends this observation by showing enhanced NLRP3 expression in BM-MNCs of AML patients and supports the assumption that aberrant kinase activity contributes to overexpression of NLRP3, since correlation analyses indicate an association between KRAS, FLT3 and NRAS with NLRP3 in AML patient BM samples. In addition, we showed that especially AML types classified as M5 (according to the FAB classification system^24^) highly express NLRP3, suggesting that monocytic AML cells that additionally carry kinase mutations might express high levels of NLRP3 and could therefore be interesting targets for treatment with NLRP3 inhibitors. Indeed, we showed that pharmacologic inhibition of NLRP3 activation with CP-456773, and also genetic deletion of NLRP3, significantly reduced the proliferation of MOLM-13, MV4-11 and HL-60 cells, whereas HeLa cells, which lack NLRP3, were not affected by the inhibitor. This confirms the specificity of the inhibitor and suggests that even cells with low NLRP3 protein expression, as shown by experiments using HL-60 cells, might be sensitive to NLRP3 inflammasome inhibition. Our further analyses revealed that treatment with CP-456773 results in growth inhibition through induction of G0/G1 phase cell-cycle arrest, along with downregulation of cyclin-dependent kinases CDK4 and CDK6 and upregulation of the CDK inhibitor p21. Dysregulation of CDK4 and CDK6 leading to uncontrolled cell-cycle progression is frequently observed in cancer.^40^ Targeting CDK4/CDK6 by selective inhibitors and prolonged arrest in the G0/G1 phase has therefore been suggested as a strategy to attenuate cell proliferation in myeloma and AML.^41–43^ Since we showed in our study that inhibition of NLRP3 activity also led to downregulation of CDK4/CDK6 and prolonged G0/G1 phase arrest, targeting NLRP3 with inhibitors might also be a promising treatment option in AML.

We further showed that MOLM-13 cells expressing high levels of NLRP3 protein efficiently engrafted and expanded in immune deficient mice, leading to marked clinical signs. In contrast, injection of MOLM-13 ΔNLRP3 cells (in which NLRP3 was genetically ablated) resulted in reduced engraftment and AML expansion, as well as fewer and less severe clinical signs, highlighting the role of NLRP3 as novel driver in AML.

Although NLRP3 was initially described in fully differentiated innate immune cells like monocytes, macrophages and dendritic cells,^2,44,45^ the protein is also expressed in human HSPCs, which are at the top of a hierarchical differentiation process and responsible for the reconstitution of all major types of blood cells.^46^ As reviewed by Ratajczak and colleagues, the NLRP3 inflammasome has multiple functions during hematopoiesis, including HSPC-expansion, mobilization, homing and engraftment as well as in aging and regulation of hematopoietic cell metabolism.^6^ Thus, it is likely that misregulation of NLRP3 inflammasomes can cause hematopoietic diseases. A better understanding of NLRP3 inflammasome activation in the context of normal and malignant hematopoiesis is therefore of great importance.

By means of shotgun proteome analysis utilizing multidimensional high-performance liquid chromatography-tandem mass spectrometry, we analyzed the potential effects of the NLRP3 inflammasome inhibitor CP-456773 in more detail. Our data suggest that inhibition of inflammasome activation results in strong regulation of the eIF2 signaling pathway. EIF2 is a key mediator of global protein synthesis. It acts as a GTP-binding protein and forms a ternary complex composed of eIF2, GTP and the methionine-loaded initiator tRNA.^47^ Phosphorylation of the eIF2α subunit hinders ternary complex formation and inhibits global protein synthesis.^30^ In addition, eIF2α phosphorylation can induce the translation of Activating Transcription Factor 4 (ATF4), which links eIF2α phosphorylation to cell cycle arrest and apoptosis.^48^ In a murine model of HER2^+^ breast cancer, eIF2α phosphorylation showed anti-tumor functions by inducing ATF4, which upregulates the CDK inhibitor p21. The latter study further suggests potential anti-tumor effects using eIF2α-phosphatase inhibitors to achieve sustained eIF2α-phosphorylation.^49^ In line with this, our study links eIF2 phosphorylation to an increase in p21, along with a decrease in CDK4, CDK6 and prolonged G0/G1 arrest, suggesting that NLRP3 inhibition blocks cell growth by inducing a p-eIF2α-dependent cell cycle arrest in the G0/G1 phase through inhibition of CDK4 and CDK6 expression and activation of p21 expression.

In conclusion, this study identifies the NLRP3/eIF2 axis as a driving force in AML. Although the exact interplay between NLRP3 and potential kinases, which contribute to the phosphorylation of eIF2α needs to be further investigated, the data presented suggest that NLRP3 inhibition might offer a new therapeutic strategy for those AML subtypes characterized by constitutive NLRP3 overexpression and/or activation.

## Acknowledgments

This work was supported by the County of Salzburg, Cancer Cluster Salzburg [grant number 20102-P1601064-FPR01-2017], the Biomed Center Salzburg (project 20102-F1901165-KZP), the European Interreg project EPIC (grant number ITAT1054), the Austrian Science Fund (FWF) [grant number P33969], and by the Priority program ACBN, University of Salzburg.

## Authorship Contribution

J.H.H., M.L., C.H. and F.A. conceptually designed experiments; M.L., L.U., T.N., P.K., R.B., H.S., D.E., H.H.D., S.R., C.B. O.B., L.Z., S.B. and S.T. acquired, analyzed, and interpreted data; D.S., R.G., L.P., T.H. and D.N. obtained patient samples; D.N. performed immunohistochemistry analyses; J.V., S.S. and S.W. performed bioinformatics and statistical analyses; P.K., D.E., S.T., M.L., H.S., R.B. and H.H.D. performed in vivo studies in NSGS mice; J.H.H., S.W., C.H. and F.A. supervised the study.

## References

1. Grivennikov SI, Greten FR, Karin M. Immunity, inflammation, and cancer. Cell. 2010;140(6):883–899.

2. Franchi L, Eigenbrod T, Munoz-Planillo R, Nunez G. The inflammasome: a caspase-1-activation platform that regulates immune responses and disease pathogenesis. Nature Immunology. 2009;10(3):241–247.

3. Kanneganti TD. The inflammasome: firing up innate immunity. Immunol Rev. 2015;265(1):1–.

4. Martinon F, Burns K, Tschopp J. The inflammasome: a molecular platform triggering activation of inflammatory caspases and processing of proIL-beta. Mol Cell. 2002;10(2):417–426.

5. Lamkanfi M. Emerging inflammasome effector mechanisms. Nat Rev Immunol. 2011;11(3):213–220.

6. Ratajczak MZ, Bujko K, Cymer M, et al. The Nlrp3 inflammasome as a “rising star” in studies of normal and malignant hematopoiesis. Leukemia. 2020;34(6):1512–1523.

7. Basiorka AA, McGraw KL, Eksioglu EA, et al. The NLRP3 inflammasome functions as a driver of the myelodysplastic syndrome phenotype. Blood. 2016;128(25):2960–2975.

8. Hamarsheh S, Osswald L, Saller BS, et al. Oncogenic Kras(G12D) causes myeloproliferation via NLRP3 inflammasome activation. Nat Commun. 2020;11(1):1659.

9. Cozzolino F, Rubartelli A, Aldinucci D, et al. Interleukin 1 as an autocrine growth factor for acute myeloid leukemia cells. Proc Natl Acad Sci U S A. 1989;86(7):2369–2373.

10. Dinarello CA. Why not treat human cancer with interleukin-1 blockade? Cancer Metastasis Rev. 2010;29(2):317–329.

11. Sanjana NE, Shalem O, Zhang F. Improved vectors and genome-wide libraries for CRISPR screening. Nat Methods. 2014;11(8):783–784.

12. Doench JG, Fusi N, Sullender M, et al. Optimized sgRNA design to maximize activity and minimize off-target effects of CRISPR-Cas9. Nat Biotechnol. 2016;34(2):184–191.

13. Sanson KR, Hanna RE, Hegde M, et al. Optimized libraries for CRISPR-Cas9 genetic screens with multiple modalities. Nat Commun. 2018;9(1):5416.

14. Kasper M, Regl G, Eichberger T, Frischauf AM, Aberger F. Efficient manipulation of Hedgehog/GLI signaling using retroviral expression systems. Methods Mol Biol. 2007;397:67–78.

15. Detre S, Saclani Jotti G, Dowsett M. A “quickscore” method for immunohistochemical semiquantitation: validation for oestrogen receptor in breast carcinomas. J Clin Pathol. 1995;48(9):876–878.

16. Dwivedi RC, Spicer V, Harder M, et al. Practical implementation of 2D HPLC scheme with accurate peptide retention prediction in both dimensions for high-throughput bottom-up proteomics. Anal Chem. 2008;80(18):7036–7042.

17. Gilar M, Olivova P, Daly AE, Gebler JC. Orthogonality of separation in two-dimensional liquid chromatography. Anal Chem. 2005;77(19):6426–6434.

18. Tyanova S, Cox J. Perseus: A Bioinformatics Platform for Integrative Analysis of Proteomics Data in Cancer Research. Methods Mol Biol. 2018;1711:133–148.

19. Cox J, Mann M. MaxQuant enables high peptide identification rates, individualized p.p.b.- range mass accuracies and proteome-wide protein quantification. Nat Biotechnol. 2008;26(12):1367–1372.

20. UniProt C. UniProt: a worldwide hub of protein knowledge. Nucleic Acids Res. 2019;47(D1):D506–D515.

21. Perez-Riverol Y, Csordas A, Bai J, et al. The PRIDE database and related tools and resources in 2019: improving support for quantification data. Nucleic Acids Res. 2019;47(D1):D442–D450.

22. Haferlach T, Kohlmann A, Wieczorek L, et al. Clinical utility of microarray-based gene expression profiling in the diagnosis and subclassification of leukemia: report from the International Microarray Innovations in Leukemia Study Group. J Clin Oncol. 2010;28(15):2529–2537.

23. Kohlmann A, Kipps TJ, Rassenti LZ, et al. An international standardization programme towards the application of gene expression profiling in routine leukaemia diagnostics: the Microarray Innovations in LEukemia study prephase. Br J Haematol. 2008;142(5):802–807.

24. Bennett JM, Catovsky D, Daniel MT, et al. Proposals for the classification of the acute leukaemias. French-American-British (FAB) co-operative group. Br J Haematol. 1976;33(4):451–458.

25. Metzeler KH, Hummel M, Bloomfield CD, et al. An 86-probe-set gene-expression signature predicts survival in cytogenetically normal acute myeloid leukemia. Blood. 2008;112(10):4193–4201.

26. Edgar R, Domrachev M, Lash AE. Gene Expression Omnibus: NCBI gene expression and hybridization array data repository. Nucleic Acids Res. 2002;30(1):207–210.

27. Coll RC, Robertson AA, Chae JJ, et al. A small-molecule inhibitor of the NLRP3 inflammasome for the treatment of inflammatory diseases. Nat Med. 2015;21(3):248–255.

28. Jiang H, He H, Chen Y, et al. Identification of a selective and direct NLRP3 inhibitor to treat inflammatory disorders. J Exp Med. 2017;214(11):3219–3238.

29. Jackson RJ, Hellen CU, Pestova TV. The mechanism of eukaryotic translation initiation and principles of its regulation. Nat Rev Mol Cell Biol. 2010;11(2):113–127.

30. Jennings MD, Kershaw CJ, Adomavicius T, Pavitt GD. Fail-safe control of translation initiation by dissociation of eIF2alpha phosphorylated ternary complexes. Elife. 2017;6.

31. Hamanaka RB, Bennett BS, Cullinan SB, Diehl JA. PERK and GCN2 contribute to eIF2alpha phosphorylation and cell cycle arrest after activation of the unfolded protein response pathway. Mol Biol Cell. 2005;16(12):5493–5501.

32. Saelens X, Kalai M, Vandenabeele P. Translation inhibition in apoptosis: caspase-dependent PKR activation and eIF2-alpha phosphorylation. J Biol Chem. 2001;276(45):41620–41628.

33. Asghar U, Witkiewicz AK, Turner NC, Knudsen ES. The history and future of targeting cyclin-dependent kinases in cancer therapy. Nat Rev Drug Discov. 2015;14(2):130–146.

34. Binder S, Luciano M, Horejs-Hoeck J. The cytokine network in acute myeloid leukemia (AML): A focus on pro- and anti-inflammatory mediators. Cytokine Growth Factor Rev. 2018;43:8–15.

35. Urwanisch L, Luciano M, Horejs-Hoeck J. The NLRP3 Inflammasome and Its Role in the Pathogenicity of Leukemia. Int J Mol Sci. 2021;22(3).

36. Hamarsheh S, Zeiser R. NLRP3 Inflammasome Activation in Cancer: A Double-Edged Sword. Front Immunol. 2020;11:1444.

37. Ju M, Bi J, Wei Q, et al. Pan-cancer analysis of NLRP3 inflammasome with potential implications in prognosis and immunotherapy in human cancer. Brief Bioinform. 2020.

38. Karki R, Kanneganti TD. Diverging inflammasome signals in tumorigenesis and potential targeting. Nat Rev Cancer. 2019;19(4):197–214.

39. Sharma BR, Kanneganti TD. NLRP3 inflammasome in cancer and metabolic diseases. Nat Immunol. 2021.

40. Malumbres M, Barbacid M. Cell cycle kinases in cancer. Curr Opin Genet Dev. 2007;17(1):60–65.

41. Huang X, Di Liberto M, Jayabalan D, et al. Prolonged early G(1) arrest by selective CDK4/CDK6 inhibition sensitizes myeloma cells to cytotoxic killing through cell cycle-coupled loss of IRF4. Blood. 2012;120(5):1095–1106.

42. Lee DJ, Zeidner JF. Cyclin-dependent kinase (CDK) 9 and 4/6 inhibitors in acute myeloid leukemia (AML): a promising therapeutic approach. Expert Opin Investig Drugs. 2019;28(11):989–1001.

43. Yang C, Boyson CA, Di Liberto M, et al. CDK4/6 Inhibitor PD 0332991 Sensitizes Acute Myeloid Leukemia to Cytarabine-Mediated Cytotoxicity. Cancer Res. 2015;75(9):1838–1845.

44. Elliott EI, Sutterwala FS. Monocytes Take Their Own Path to IL-1 beta. Immunity. 2016;44(4):713–715.

45. Netea MG, Nold-Petry CA, Nold MF, et al. Differential requirement for the activation of the inflammasome for processing and release of IL-1 beta in monocytes and macrophages. Blood. 2009;113(10):2324–2335.

46. Lenkiewicz AM, Adamiak M, Thapa A, et al. The Nlrp3 Inflammasome Orchestrates Mobilization of Bone Marrow-Residing Stem Cells into Peripheral Blood. Stem Cell Reviews and Reports. 2019;15(3):391–403.

47. Merrick WC, Pavitt GD. Protein Synthesis Initiation in Eukaryotic Cells. Cold Spring Harb Perspect Biol. 2018;10(12).

48. Wortel IMN, van der Meer LT, Kilberg MS, van Leeuwen FN. Surviving Stress: Modulation of ATF4-Mediated Stress Responses in Normal and Malignant Cells. Trends Endocrinol Metab. 2017;28(11):794–806.

49. Darini C, Ghaddar N, Chabot C, et al. An integrated stress response via PKR suppresses HER2+ cancers and improves trastuzumab therapy. Nat Commun. 2019;10(1):2139.

