## Supplemental data for "The NLRP3/eIF2 axis drives cell cycle progression in acute myeloid leukemia"

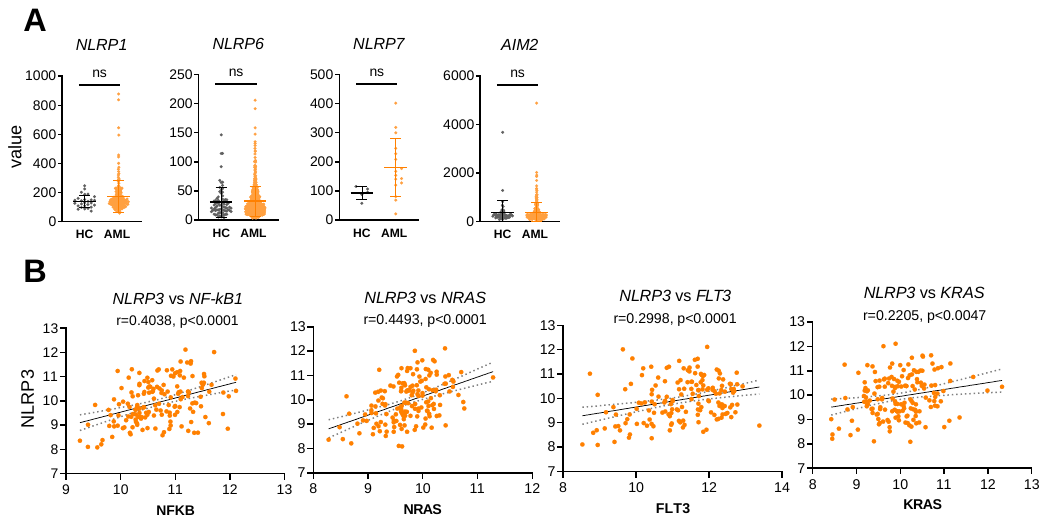

**Supplemental Figure 1**

**Expression of NLR family members in AML.** (A) NLRP1, NLRP6, NLRP7 and AIM2 expression in AML patients (AML, n=542) compared to healthy controls (HC, n=74) was determined from publicly available dataset GSE13159. The dataset was analyzed using Python. A two-tailed, unpaired t test was performed for the statistical analysis. Single dots represent individual donors; the horizontal lines in each column represent mean ± SD. (B) Correlation analysis between NLRP3, NF-κB and NRAS, FLT3 and KRAS in AML patients using the dataset GSE12417 (n=163). The GEOparse package was used for data access and Python for data processing and statistical analysis.

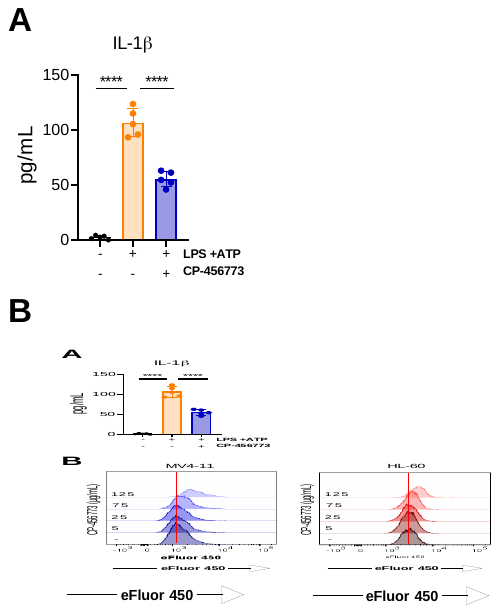

**Supplemental Figure 2.**

**The NLRP3 inhibitor CP-456773 inhibits the release of IL-1β and suppresses the proliferation of NLRP3-expressing AML cell lines.** (A) MOLM-13 cells were seeded and stimulated with E. coli lipopolysaccharide (LPS, 50 ng/ml) for 4 hours and then treated with CP-456773. After 2 hours of NLRP3 inhibition, cells were stimulated with adenosine triphosphate (ATP, 5 mM) for 16 hours to induce NLRP3 inflammasome activation. Supernatants were removed and analyzed for IL-1β secretion by ELISA. Individual dots represent single experiments, bars represent mean ± SD. (B) Proliferation of MV4-11 and HL-60 cells in the presence of the indicated concentrations of CP-456673 was monitored after 72 hours by flow cytometry using the Cell Proliferation Dye eFluor 450. Histograms of one representative out of 5 experiments are shown. The vertical red lines indicate the fluorescence peak of proliferating cells in the absence of CP-456773.

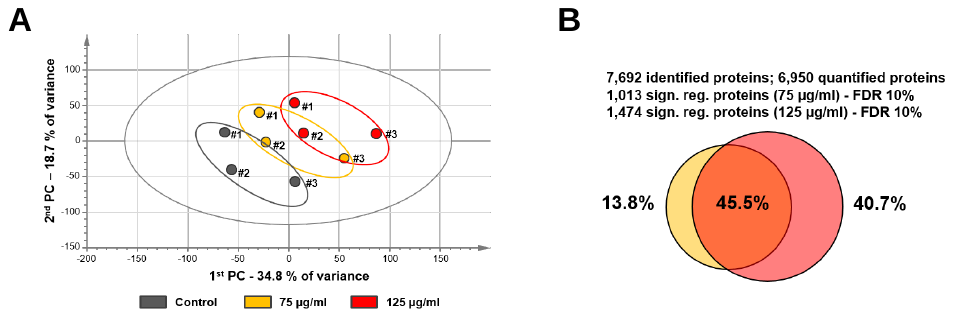

**Supplemental Figure 3.**

**Differential proteomic analyses.** (A) Multivariate statistics employing principal component analysis (PCA) reveals grouping of the three different treatments according to their shift in detected protein expression. Expression values of 6,950 proteins that were quantified in each of the samples were used to generate the plot. (B) A Venn diagram depicts the overlap (orange) between significantly regulated proteins (FDR 10%) after treatment with 75 µg/mL CP-456773 (yellow) and 125 µg/mL CP-456773 (red).

**Proteomic analysis of MOLM-13 – Supplemental version**

**Chemicals.** Dithiothreitol (DTT, ≥ 99.5%), formic acid (FA, 98.0-100%), iodoacetamide (IAA, ≥ 99.0%), sodium dodecyl sulfate (SDS, ≥ 99.5%) and triethylammonium bicarbonate (TEAB, pH 8.5, 1 mol/L were obtained from Sigma-Aldrich (Vienna, Austria). Acetonitrile (ACN, ≥ 99.9%) and methanol (MeOH, ≥ 99.9%) were obtained from VWR International (Vienna, Austria). Ammonia (25%) and ortho-phosphoric acid (85%) were purchased from Merck (Burlington, MA, USA). Trypsin (sequencing grade modified, porcine) was obtained from Promega (Madison, WI, USA). A MilliQ Integral 3 instrument (Millipore, Billerica, MA, USA) was used for deionization of water.

**Cell culture.** MOLM-13 cells were seeded in a 48-well plate at a density of 2×10^5^/mL (1×10^5^/500 µL/well). Cells were treated with 75 and 125 µg/mL of CP-456773 and incubated for 24 h at 37°C and 5% CO2. DMSO-treated cells were used as a control. Biological replicates were generated by conducting this treatment scheme on three consecutive days. Cells were washed thoroughly with PBS before sample preparation.

**Sample preparation.** S-Trap mini columns (Protifi, Huntington, NY, USA) were employed according to the manufacturer´s instructions with minor adjustments: a cell pellet of approximately 1×10^6^ cells was lysed in 5% SDS and 50 mmol/L TEAB (pH 7.55) at 95°C for 5 min followed by sonication in a Bioruptor device (Diagenode, Liège, Belgium) for 10 min. After a centrifugation step, protein content was analyzed by a Pierce BCA Protein assay kit (Thermo Fisher Scientific, Vienna, Austria). Denaturation and reduction of proteins were performed by supplementation of DTT to 40 mmol/L and incubation at 95°C for 10 min. Reduced cysteine residues were alkylated by the addition of IAA to a concentration 80 mmol/and incubation at 21°C in the dark for 30 min. After a precipitation step and thorough washing, 10 µg of trypsin were solubilized in 50 mmol/L TEAB (pH 8.5), added to the S-Trap matrix and incubated at 37°C for 18 h. Peptides were eluted and subsequently dried at 30°C using a vacuum centrifuge. Samples were resuspended in 100 mmol/L TEAB (pH 8.5) to a concentration of 1.0 mg/mL. 100 µg of peptides of each sample were labeled by using a TMT 10plex™ kit (Thermo Fisher Scientific) according to the manufacturer´s instructions. Samples were labeled with the tags 126 to 130C (Figure 4). All nine samples were pooled and dried at 30°C in a vacuum centrifuge. The combined sample was resuspended in H_2_O + 20 mmol/L ammonium formate (pH 10.0) to a concentration of 1.8 mg/mL.

**High-performance liquid chromatography and mass spectrometry.** The pooled peptide sample was subjected to high-pH reversed phase fractionation without any prior purification. This off-line fractionation was carried out on an Agilent 1100 Series capillary LC system (Santa Clara, CA, USA). The peptides were separated on two sequentially linked Gemini NX-C18 columns (150 × 2.0 mm i.d., 3 µm particle diameter, 110 Å pore size) purchased from Phenomenex Inc. (Aschaffenburg, Germany) that were connected by a short 20 µm i.d. connective tubing. Mobile phases A (H2O + 20 mmol/L ammonium formate, pH 10) and B [(90.0% ACN / 10.0% H2O) + 20 mmol/L ammonium formate, pH 10] were prepared according to Dwivedi et al. and Gilar et al. [1,2]. A stepped linear gradient was applied: 1.0% B for 15.0 min, 1.0% - 30.0% B for 185.0 min, 30.0% - 60.0% for 30.0 min, 80.0% B for 30.0 min and 1.0% B for 40.0 min. The flow rate was set to 150 µL/min, the column oven temperature to 40°C, and 100.0 µl of sample were injected. 30 independent fractions were collected by an integrated automatic sample fraction collector system at uniform time slices starting at time point 12.0 min until time point 264.0 min. These 30 fractions were pooled into six fractions employing sample concatenation as reviewed by Yang et al., increasing the orthogonality of the two chromatographic dimensions both relying on reversed phase separation principles [3]. Pooled fractions were dried at 45°C in a vacuum centrifuge and resuspended in 16 µl of H2O + 0.1% FA.

As a second dimension, acidic reversed phase HPLC was employed using a 2000 mm µPAC™ C18 column (PharmaFluidics, Ghent, Belgium). These nanoscale chromatographic separations were carried out on a nanoHPLC instrument (UltiMate™ U3000 RSLCnano, Thermo Fisher Scientific, Germering, Germany) at a flow rate of 300 nL/min and a column oven temperature of 50°C. Mobile phase solution A contained H2O + 0.10% FA, and mobile phase B contained ACN + 0.10% FA. After solvent B was kept at 1.0% B for 5.0 min, a linear gradient to 40.0% B in 595.0 min was applied. This gradient was followed by a purging step at 90.0% B for 30.0 min. The column was re-equilibrated at 1.0% B for 100.0 min. 1.0 µl of each fraction was injected using a microliter pick-up mode (5.0 µl loop volume). Each fraction was measured once.

The nanoHPLC was hyphenated to a quadrupole-Orbitrap hybrid mass spectrometer (Thermo Scientific QExactive Plus benchtop quadrupole-Orbitrap mass spectrometer) via a Nanospray Flex ion source (both from Thermo Fisher Scientific, Bremen, Germany). The source was equipped with a SilicaTip emitter with 360 µm o.d., 20 µm i.d. and a tip i.d. of 10 µm purchased from New Objective (Woburn, MA, USA). The mass spectrometer was operated with the following instrument settings: spray voltage of 1.5 kV, S-lens RF level of 55.0, capillary temperature of 320 °C and an MS1 AGC target of 3e6 in an m/z range of 400-2000 with a maximum injection time of 100 ms. A MS1 scan at a resolution setting of 70,000 at 200 m/z was followed by 15 data-dependent MS2 scans at a resolution of 35,000 at m/z 200. Target peptides were fragmented by HCD at 32.0 NCE in a 2.0 m/z isolation window with an AGC target of 1e5 and a maximum injection time of 100 ms. A dynamic exclusion setting of 30 s was applied. Pierce LTQ Velos ESI Positive Ion Calibration Solution from Life Technologies (Vienna, Austria) was used for calibration of the instrument.

**Data Evaluation.** Acquired raw data were evaluated using MaxQuant software [4] in default settings correcting for isotope impurities in TMT reagents (provided by the manufacturer). Uniprot database entries including both Swiss-Prot and TrEMBL entries for *Homo sapiens* (access: 10.03.2019) were provided for MaxQuant protein identification [5]. The obtained protein groups were further processed using Perseus: Protein groups were filtered removing potential contaminants, proteins that were only identified by site and reverse sequence matches [6]. Only those protein groups providing quantitative values for all nine reporter ion channels were further processed, including log2-transformation and normalization by subtraction of the median. For statistical analysis and data representation, R-software version 3.6.1 as well as GraphPad Prism version 8.0.2. (GraphPad Software, San Diego, CA, USA) were employed [7]. Venn diagrams were compiled with the Venn diagram plotter developed by the Pacific Northwest National Laboratory (PNNL) (https://omics.pnl.gov/software/venn-diagram-plotter). Principal component analysis was conducted utilizing Simca 13.0.3 (Umetrics, Sartorius Stedim Biotech, Göttingen, Germany) using unit variances and mean centering as data pre-processing steps. For pathway analysis, significantly regulated proteins (Benjamini-Hochberg corrected p-value of ≤0.1) were used as input for Ingenuity Pathway Analysis (Qiagen, CA, USA).

**Inflammasome activation assay and validation of the NLRP3 inhibitor.** CP-456773 sodium salt ≥98% (HPLC) (Sigma) was dissolved according to the manufacturer’s instructions. MOLM-13 cell lines were seeded in a 48-well plate at a density of 2×10^5^/mL (1×10^5^/500 µL/well). To assess the functionality of CP-456773, the cells were stimulated with 50 ng/mL LPS from *Escherichia coli* 055:B5 (Sigma) for 4 h and treated with the NLRP3 inhibitor. Cells were then stimulated with 5 mM adenosine 5′-triphosphate disodium salt hydrate (ATP) for 16 h (InvivoGen). Supernatants were removed and analyzed for IL-1β using commercially available ELISA kits (R&D Systems), according to the manufacturer’s instructions.

1.	Gilar, M.; Olivova, P.; Daly, A.E.; Gebler, J.C. Orthogonality of separation in two-dimensional liquid chromatography. *Anal Chem* **2005**, *77*, 6426-6434, doi:10.1021/ac050923i.

2.	Dwivedi, R.C.; Spicer, V.; Harder, M.; Antonovici, M.; Ens, W.; Standing, K.G.; Wilkins, J.A.; Krokhin, O.V. Practical implementation of 2D HPLC scheme with accurate peptide retention prediction in both dimensions for high-throughput bottom-up proteomics. *Anal Chem* **2008**, *80*, 7036-7042, doi:10.1021/ac800984n.

3.	Yang, F.; Shen, Y.; Camp, D.G., 2nd; Smith, R.D. High-pH reversed-phase chromatography with fraction concatenation for 2D proteomic analysis. *Expert Rev Proteomics* **2012**, *9*, 129-134, doi:10.1586/epr.12.15.

4.	Cox, J.; Mann, M. MaxQuant enables high peptide identification rates, individualized p.p.b.-range mass accuracies and proteome-wide protein quantification. *Nat Biotechnol* **2008**, *26*, 1367-1372, doi:10.1038/nbt.1511.

5.	UniProt, C. UniProt: a worldwide hub of protein knowledge. *Nucleic Acids Res* **2019**, *47*, D506-D515, doi:10.1093/nar/gky1049.

6.	Tyanova, S.; Cox, J. Perseus: A Bioinformatics Platform for Integrative Analysis of Proteomics Data in Cancer Research. *Methods Mol Biol* **2018**, *1711*, 133-148, doi:10.1007/978-1-4939-7493-1_7.

7.	R-Core-Team. R: A language and environment for statistical computing. *R Foundation for Statistical Computing, Vienna, Austria* **2013**.
